# MLDAAPP: Machine Learning Data Acquisition for Assessing Population Phenotypes

**DOI:** 10.1101/2023.09.20.558733

**Authors:** Amir R. Gabidulin, Seth M. Rudman

## Abstract

Collecting phenotypic data from many individuals is critical to numerous biological disciplines. Yet, organismal phenotypic or trait data are still often collected manually, limiting the scale of data collection, precluding reproducible workflows, and creating the potential for human bias. Computer vision could largely ameliorate these issues, but currently available packages only operate with specific inputs and hence are not scalable or accessible for many biologists. We present Machine Learning Data Acquisition for Assessing Population Phenotypes (MLDAAPP), a package of tools for collecting phenotypic data from groups of individuals. We demonstrate that MLDAAPP is both accurate and uniquely effective at measuring phenotypes in challenging conditions - particularly images and videos of varying quality derived from both lab and field environments. Employing MLDAAPP solves key issues of reproducibility, increases both the scale and scope of data generation, and reduces the potential for human bias.

## Introduction

Biological measurements on groups of individuals, including counts, sizes, and patterns of movement, are essential data across a range of disciplines including ecology, evolutionary biology, animal behavior, and genetics ^1–4^. Collection of these data are still primarily done manually, which limits both the scope and reproducibility of the resultant outputs ^5–7^. The rapid growth of Machine Learning (ML) is potentially transformative to the way data is collected as demonstrated by successful implementations in health sciences ^8,9^. The automated extraction of information from images and videos, often called computer vision (CV), is a particularly promising area for streamlining biological data generation ^7,10^. Existing ML programs present considerable barriers to the collection of whole organism phenotypic data. These include monetary cost, implementation requiring considerable coding expertise, and designs optimized for the collection of data only under very specific circumstances ^7,11^. These challenges can be insurmountable when researchers seek to use CV methods to collect phenotypic data in non-standard assays, field research environments, on groups of organisms, or any combination of these domains ^6,11,12^.

Applications of CV have successfully generated datasets for use in ecology, evolution, and genetics ^7,12–14^. These methods employ pre-defined background subtraction to allow for a streamlined approach to analyzing photo and video data. However, CV can be done with supervised machine learning approaches in which annotated training datasets are analyzed, and weighted decisions are used to create bounding boxes or instance segmentation masks around objects. Supervised machine learning approaches allow users to define what to detect rather than using an *a priori* dictated set of rules as is found in pure background subtraction approaches. This increased flexibility is tremendously beneficial if users seek to employ CV techniques to generate phenotypic data beyond laboratory environments or to use images or videos without optimized contrast ^6^.

Here, we introduce MLDAAPP, a package of scripts that leverages existing ML tools for the collection of phenotypic data focused on user simplicity, flexibility, and precision. MLDAAPP utilizes an open-source ML product YOLOv8, which uses enterprise level supervised machine learning and computer vision algorithms and offers state-of-the-art performance in speed and accuracy ^15^. The purpose of this package is to translate YOLOv8 to quantify photos and videos to produce object counts, object sizes, positions, avoidance/preference behaviors, and locomotor activity with a focus on data outputs that are simple to analyze for biologists. We combine several core components of YOLOv8’s functionality with streamlined annotation methods through Roboflow to build a workflow for a wide range of scenarios focused on the study of populations ranging from standardized environments and small numbers of individuals to complex natural environments with dynamic group compositions. In contrast with the existing ML software used to generate whole organism phenotype data, implementations of MLDAAPP require little to no investment in specialized imaging hardware or fixed computing resources - cell phone videos and low-cost cloud computing is often sufficient for accurate high-throughput data collection. MLDAAPP allows users to measure behaviors in natural conditions far beyond the shadowless high-contrast scenarios required for many background-subtraction based CV packages ^7,12^. Moreover, MLDAAPP allows for a full interactive ML model training pathway through user generated annotations, which provides a greatly increased flexibility for user image and video inputs.

We report results from generated datasets that span a wide range of phenotypes and taxa using MLDAAPP. We provide guidance on best practices and information to increase usability when employing the accompanying GitHub package^16^.We also compare the performance of MLDAAPP to both human counted data and existing programs that employ ML for the generation of group phenotypic data across a range of data collection scenarios (idtrackerai and TRex ^12,13^). The significant increase in performance and flexibility of MLDAAPP reinforce its utility to solving key issues of reproducibility ^17^, increasing the scale and scope of data generation ^2^, and greatly reducing the potential for human bias ^18^ across a range of biological disciplines.

## Methods

### Input preparation, model training, and model implementation

A complete user guide that includes information on input generation, annotation, model training, and model implementation for MLDAAPP is available on the associated GitHub^16^. Lurig et al. ^7^ provide both a history of CV for the collection of phenotypic data for biology and general best practices for input preparation that largely apply to users of MLDAAPP. Here we provide basic background information and details specific to the application of MLDAAPP.

Crucial to the implementation of MLDAAPP is the preparation of the inputs, either photos or videos, which are used to train the custom detection model. A strong training set will span both the range and the size of the inputs to be analyzed and will be properly annotated by the user. A web-based annotating service called Roboflow can significantly reduce the time required to annotate training sets at little to no cost. Across test cases, models built with 70/30 and 80/20 validation/test ratios of training inputs both yielded near identical Mean Average Precision metrics when working within challenging sample datasets (Figure S1). Once training sets have been generated it is important to optimize the given baseline models according to your needs, so the user should give special attention to various conditional syntaxes provided by YOLOv8.

MLDAAPP can be used to translate outputs from YOLOv8 for empirical research for object detection, instance segmentation, pose/keypoint estimation, and classification (Figure 1 and Figures S3-S6). Models range in size, with trade-offs between computing power and precision, ranging between YOLOv8n (nano-model) to YOLOv8x (extra-large-model). Table S2 briefly summarizes the most successful strategies in working with YOLOv8 and Botsort Tracker, an ML-based tracking algorithm that was used as an assistant in tracking objects throughout the varying trials.

**Figure 1:**
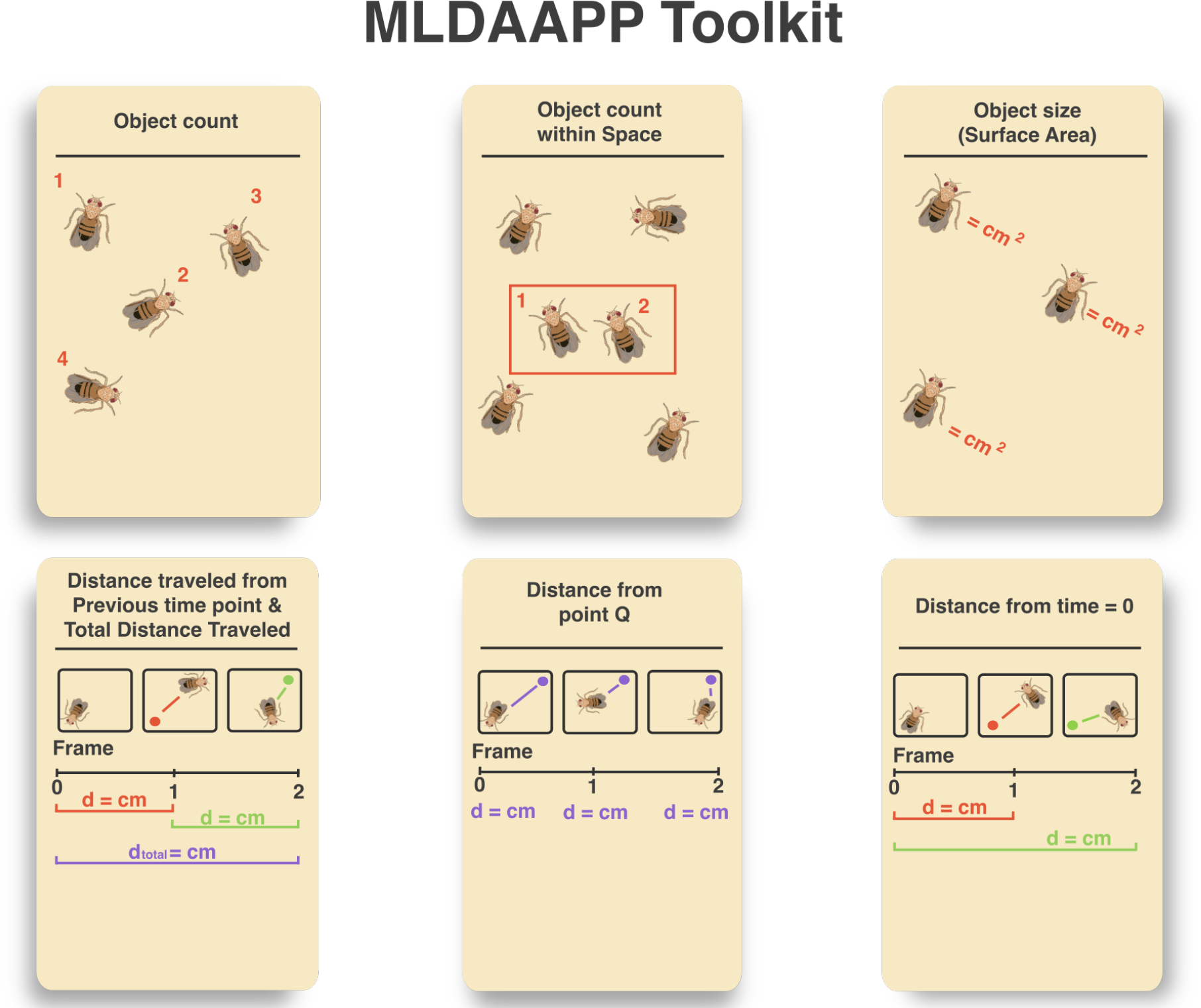
Visual demonstrations of key outputs that can be generated using user-trained computer vision models in MLDAAPP. Additional utilities can be found on the GitHub page.

## Results

### Data Acquisition and Interpretation of examples

To test MLDAAPP we both generated novel inputs across a range of organisms and sought out videos that provide realistic movement from existing sources. Below, we provide case studies on implementing three widely applicable ML data acquisition procedures, including an explicit comparison of object tracking using both MLDAAPP and idtracker.ai ^12^.

#### Object counting Case #1

Counting the number of organisms in a heterogeneous environment is time consuming and challenging, particularly when they are at high density. One common challenge in the workflow of biological investigations with the model system *Drosophila* is the estimation of the number of eggs produced (i.e., fecundity), which is often measured on large numbers of individuals resulting in an extensive expenditure of manual work hours ^19–21^. The challenge of doing this efficiently and accurately has led to the development of a variety of methods, most of which require a large time-investment into the preparation process prior to imaging ^22,23^. Here, we use MLDAAPP to automate the process of estimating fecundity of *Drosophila melanogaster* (Figure 1). We generated a dataset of 210 unique images of 35mm petri dishes in which 5 females had been allowed to lay eggs for a 24 hour period. We manually annotated images in the RoboFlow user interface to create a ML model. In cases where the input dataset is small, it is possible to enhance the training variation using Roboflow augmentation methods (see Figure S2 for details on the efficacy of this approach). Additionally, we used a “mosaic” digital augmentation as provided by Roboflow, bolstering our data set to 504 images, and training on an 88-13% split for 100 epochs. We plot the results of employing this model compared against human counts done using a microscope in Figure 2A. The model accuracy overall is high (R^2^ = 0.91), even with inputs obtained using a simple imaging setup that produces some shadows and reflection (Figure 2C).

**Figure 2A.**
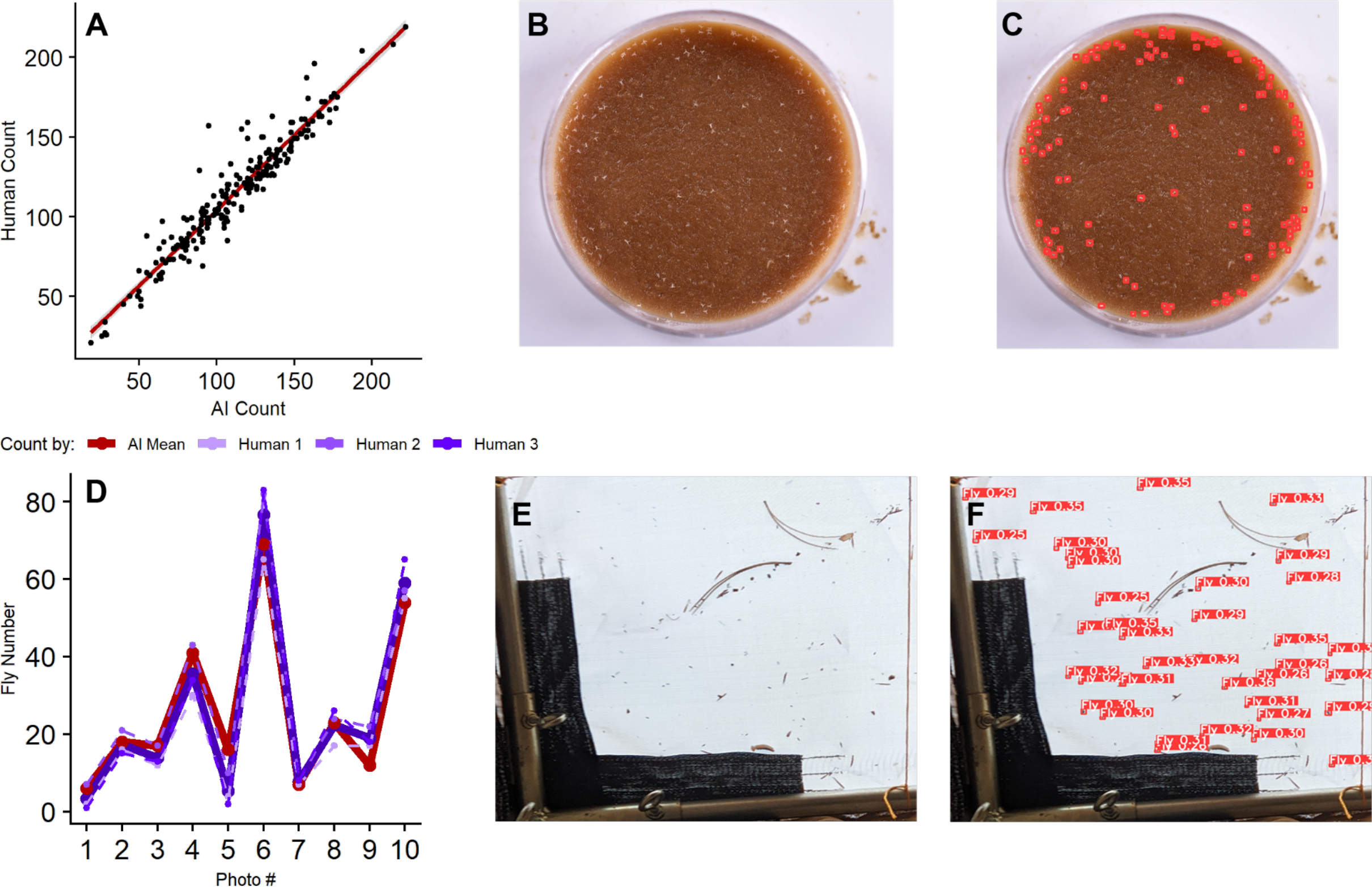
*Drosophila melanogaster* egg counts as measured by MLDAAPP vs. Human counts. Line shows linear regression of this relationship with confidence interval. Panels **B** and **C** show a representative image from a *D. melanogaster* fecundity trial before and after annotation with MLDAAPP (respectively). **D**. *Drosophila melanogaster* adult counts from outdoor population census photos as determined by MLDAAPP and three different people. MLDAAP counts are in red and mean human counts are in blue with counts from each individual represented by a dotted line. Panels **E** and **F** show a representative image from an area of an outdoor experimental cage with *D. melanogaster* before and after annotation with MLDAAPP (respectively).

#### Object Counting Case #2

Machine learning methods that employ custom annotations are also flexible enough to be used in natural environments. Hence, another application that we have developed was to provide census counts of adult *Drosophila* in large outdoor mesocosms ^20,24^ (Figure 2D). The model was trained with a data set of 80 photos using 70-30% split for 200 epochs (the passing of training data through the algorithm), where no physical or digital augmentations were used to enhance the training set . Figure 2D indicates the accuracy of the AI model given 10 census pictures in comparison to 3 independent human counts. AI model counts were positively correlated with Human 1 (r_8_ = 0.97, p < 0.001), Human 2 (r_8_ = 0.98, p < 0.001), and Human 3 (r_8_ = 0.97, p < 0.001) despite considerable image complexity (Figure 2F).

#### Object tracking Case #3

In addition to photo analysis, MLDAAPP can employ YOLOv8 with a custom tracker to generate empirical data from videos (Figure 1). We used 2 simplified standardized environments alongside 2 complex non-standardized videos to examine the flexibility of MLDAAPP for video analysis. Additionally, we analyzed two videos from the web to further demonstrate the breadth over which MLDAAPP can be used (all videos available in Table S1). Implementation was challenging in videos that had low resolution, low framerate, and small object size (Table S1 *“Daphnia”*). These conditions initially rendered object tracking unreliable as individual IDs were frequently re-assigned. This problem was largely solved by artificially increasing the number of Frames Per Second (FPS) which allowed for smoother object tracking with fewer object losses but maintaining a high FPS when filming fast objects is preferrable. MLDAAPP is effective in generating empirical data from videos with various challenges including: small and large object sizes, small and large object quantities, slow and fast object movement, low and high resolution, low and high frame rate, and simple or complex backgrounds (Table S1).

#### Comparison between MLDAAP and existing ML algorithms with similar aims

We compared the performance of MLDAAPP to idTracker.ai and TRex (Romero-Ferrero et al., 2019; Walter & Couzin, 2021), existing ML programs designed to track behavior of multiple individuals. Comparisons were made across three scenarios. First, we examined the performance of each model in tracking the movement of *Drosophila* in optimized conditions in a video provided with idTracker.ai, which was generated using a professional lighting and camera setup (>$10k in 2023). In these ideal conditions, all three algorithms successfully tracked individuals with near 100% accuracy. We next compared performance on *Drosophila* videos collected with a DSLR and lighting setup of lower cost (∼$700 in 2023) and quality which produced videos with inconsistent shading and visual noise. MLDAAPP tracked all individuals but neither idTracker.ai nor TRex were capable of consistently tracking over half of the 17 *Drosophila* and both produced false positives (Figure 3A). We next used 2 videos (Table S1 ‘Hamsters’ & ‘Honeybees’) generated by other parties that contained visual noise from shadows, inconsistent lighting, and object occlusion. Tracking performance of idTracker.ai and TRex was poor for the ‘hamsters’ video with more false positives than individuals accurately identified and tracked (Table 1). In addition, both algorithms require a static number of objects in input videos and hence were unable to run the ‘honeybees’ video due to dynamic movement of objects in and out of frame.

**Table 1.**
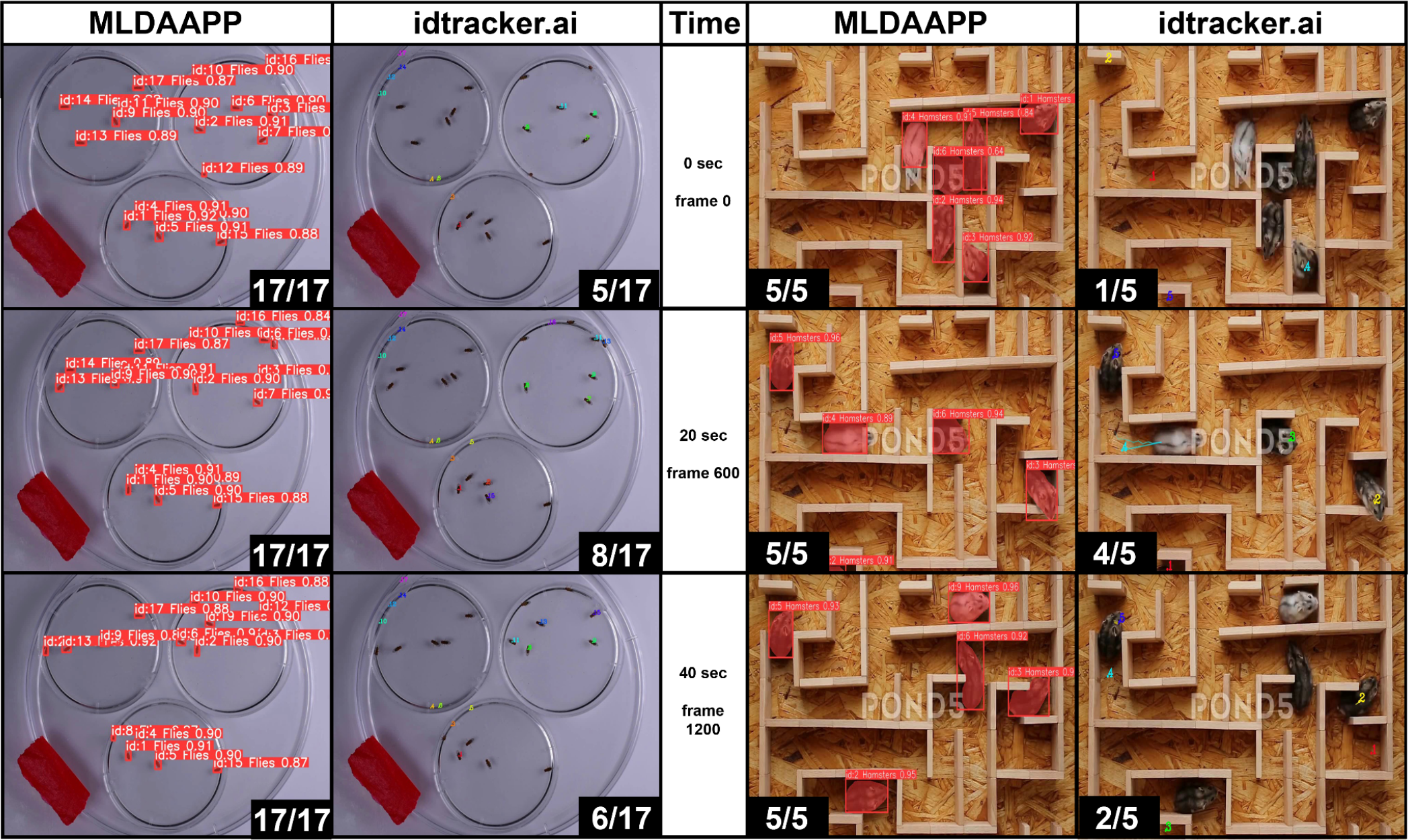
Model Comparison between MLDAAPP and idtracker.ai using non-standardized environments (Table S1). Left side shows sequential video frames tracking *Drosophila* activity with all individuals tracked in MLDAAPP and consistently >50% of individuals tracked in idtracker.ai. Right side shows results of tracking hamsters in a maze with MLDAAPP, again showing superior tracking. These videos were also conducted using TRex ^13^, which performed similarly to idtracker.ai.

Generating accurate data with background complexity is a major focus of MLDAAPP. Relaxing the need for perfect video inputs tremendously increases usability across a range of organisms and settings. In short, MLDAAPP provides a significant increase in performance in all but the most optimized conditions, providing users with the flexibility to measure phenotypes in a range of realistic settings without having to perfect imaging conditions.

## Discussion

The streamlined collection of unbiased phenotypic data is a limiting factor in many disciplines of biology ^1,2,18^. The combination of machine learning and artificial intelligence holds considerable promise for the study of a wide range of organismal traits and the speed of innovation in this area is considerable ^25^. Existing methods that use machine learning, work well for inputs that are largely standardized ranging from cells to individuals ^7,9,26^. By applying exceptional computer vision and machine learning tools, including YOLOv8, the MLDAAPP package is tractable with imperfect image and video inputs, affordable, simple to use, and built on user-generated training datasets. While we will continue to add functionality that enhances utility, MLDAAPP is a uniquely capable machine learning program for translating visual inputs to a wide range of data that are central to the study of populations.

## Notes

https://github.com/ganamir/MLDAAPP/tree/main

### Competing Interest Statement

The authors have declared no competing interest.

