## Supplemental Material for "MLDAAPP: Machine Learning Data Acquisition for Assessing Population Phenotypes"

**Supplementary Information:**

The GitHub associated with MLDAAPP contains line-by-line instructions on how to go from raw inputs (photos or videos), to custom annotations, and empirical outputs. Table S2 provides some guidelines to optimize MLDAAPP performance

The supplementary information contains:

Figure S1: Effect of training set ratio on mean Average Precision
Figure S2: Effect of training set augmentation on model accuracy
Figure S3: Total object displacement as calculated by MLDAAPP
Figure S4: Object movement from a designated point as calculated by MLDAAPP
Figure S5: Object surface area as calculated by MLDAAPP
Figure S6: Object frame-by-frame movement as calculated by MLDAAPP
Table S1: Links to images and videos used for data generation in this paper
Table S2: Suggested practices for running MLDAAPP

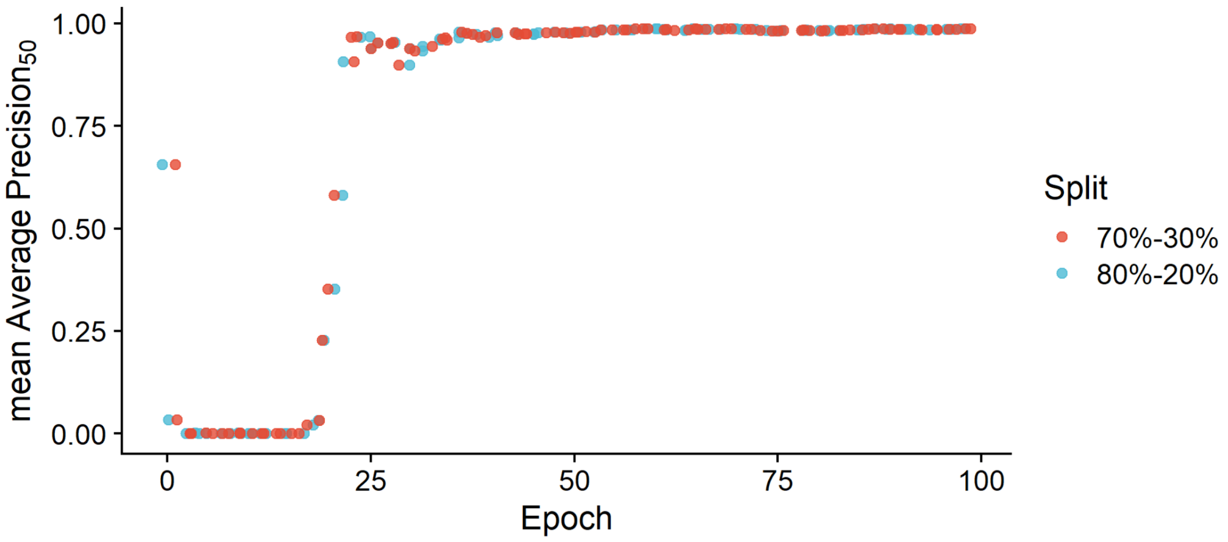


**Figure S1:** Using a Hamster data set (see table S1 for link) we compared the performance of training sets with 70/30 & 80/20 ratio splits of training-verification data on the YOLOv8x-seg model. We plot generational time (epochs) over mean Average Precision 50 (mAP50), which is a metric used to compare the model performances of object detection and segmentation systems. Here we find little difference in performance using a medium sized training set of 145 images over a generational period of 100 epochs.

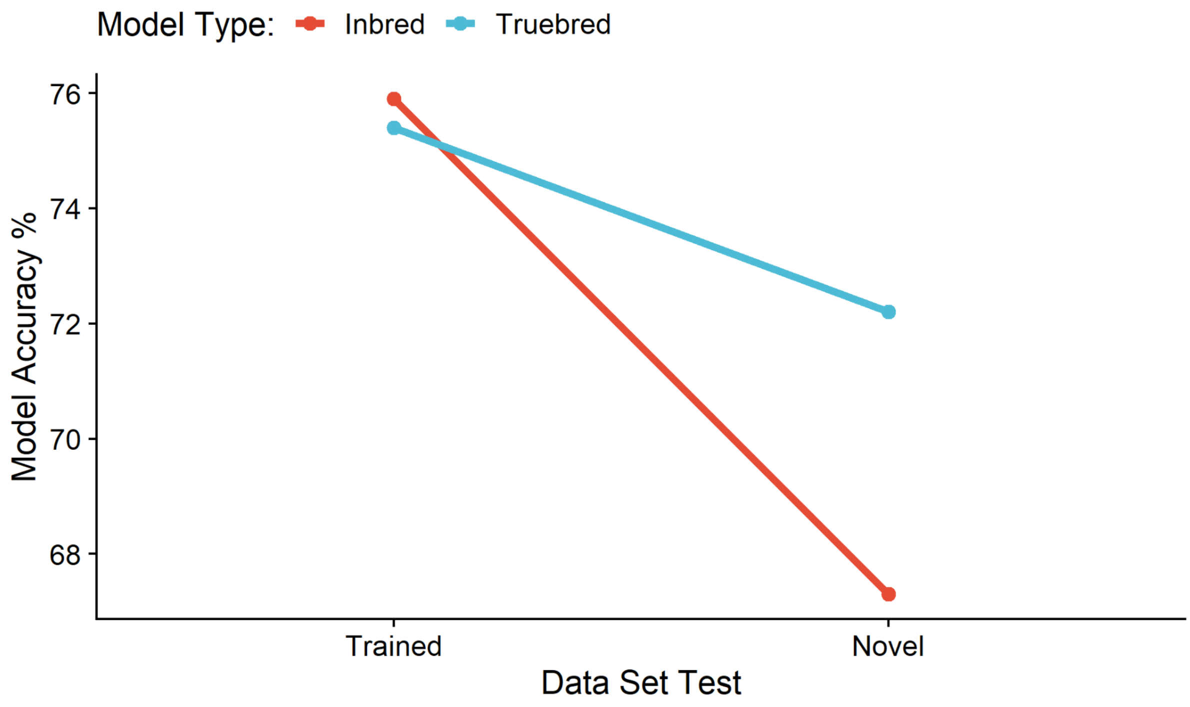


**Figure S2**: Using a fecundity trial set (see table S1 for link), we assessed performance differences of training input augmentations. The inbred model used is an augmented data-set containing 367 images constructed from 80 unique inputs. Each of the unique inputs within the data-set were randomly picked 4-5 times and both digital and physical methods were used to produce variation including camera elevation, lighting, exposure time, and background composition. A truebred model consisted of 80 unique scenarios without physically or digitally augmenting or bolstering the data-set, maintaining it at an n = 80. Both models were trained at a 70/30 ratio split for 200 epochs. The graph demonstrates the relative performance of the models compared to the human counts on the training and novel data-sets, where the novel data-set is defined as an image-set on which the models were not trained on. This graph indicates that an inbred model has been "overfitted," and therefore performs worse under "real" conditions rather than the trubred model.

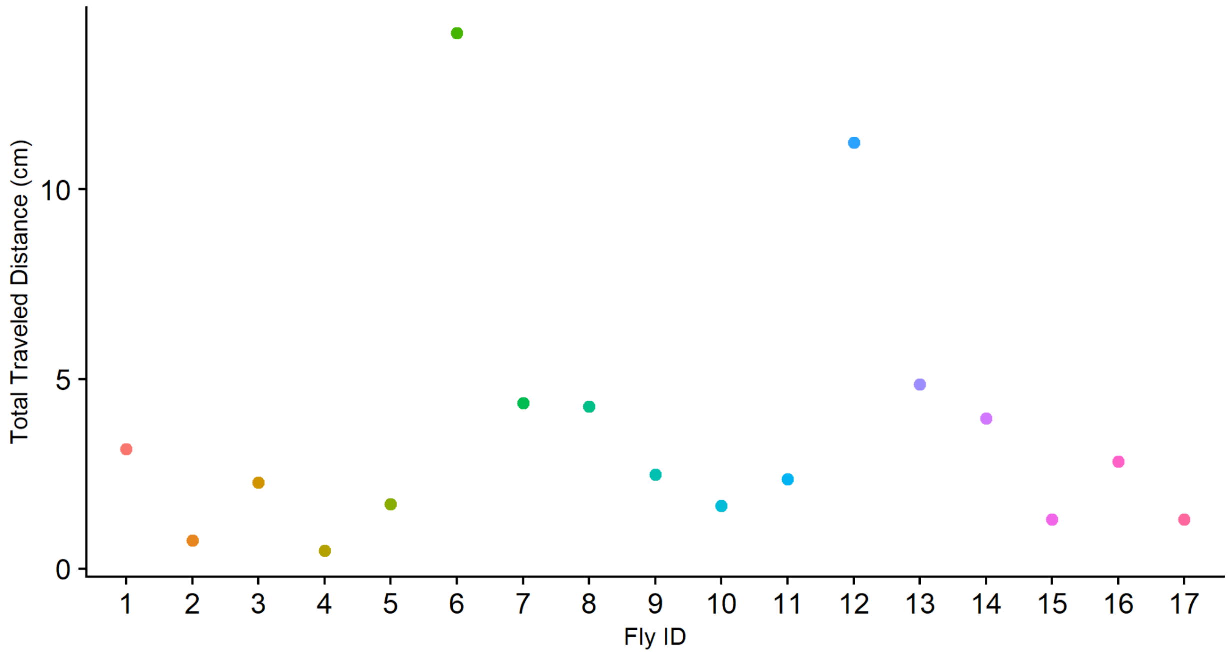


**Figure S3:** Total centroid distance displacement as calculated by summing the absolute value of displacement between frames over the entire video by individual ID using an algorithm of: n = 297, 70-30% split, 200 epochs, 60 seconds (1800 frames) of a non-standardized fruit fly videos (link in table S1).

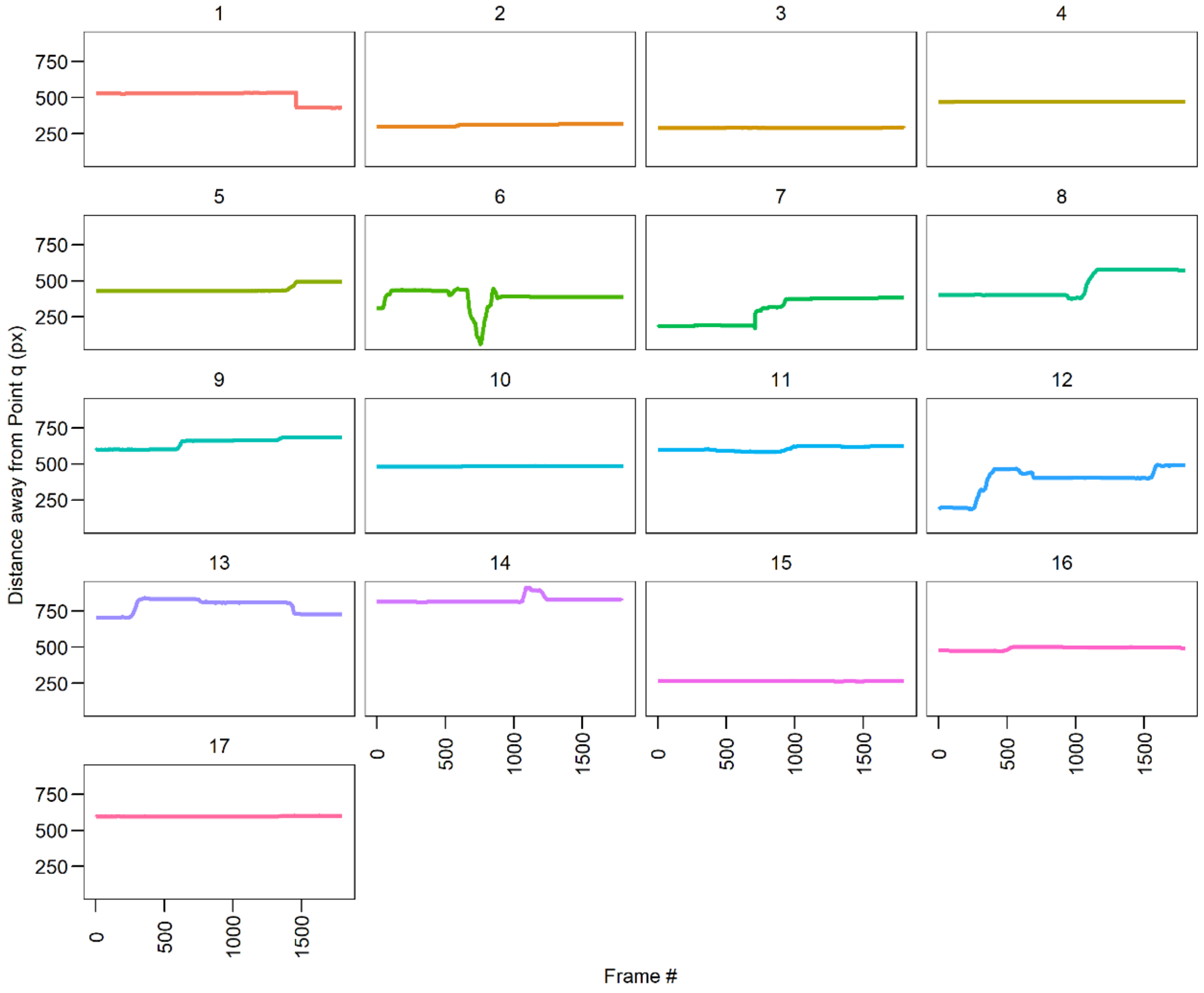


**Figure S4:** Centroid distance away from an assigned point q by individual ID in pixels using an algorithm trained on: n = 297, 70-30% split, 200 epochs, point q being a custom coordinate of space within the media as set by the user. Data extracted using a 60 second (1800 frames) non-standardized fruit fly environment (link in table S1).

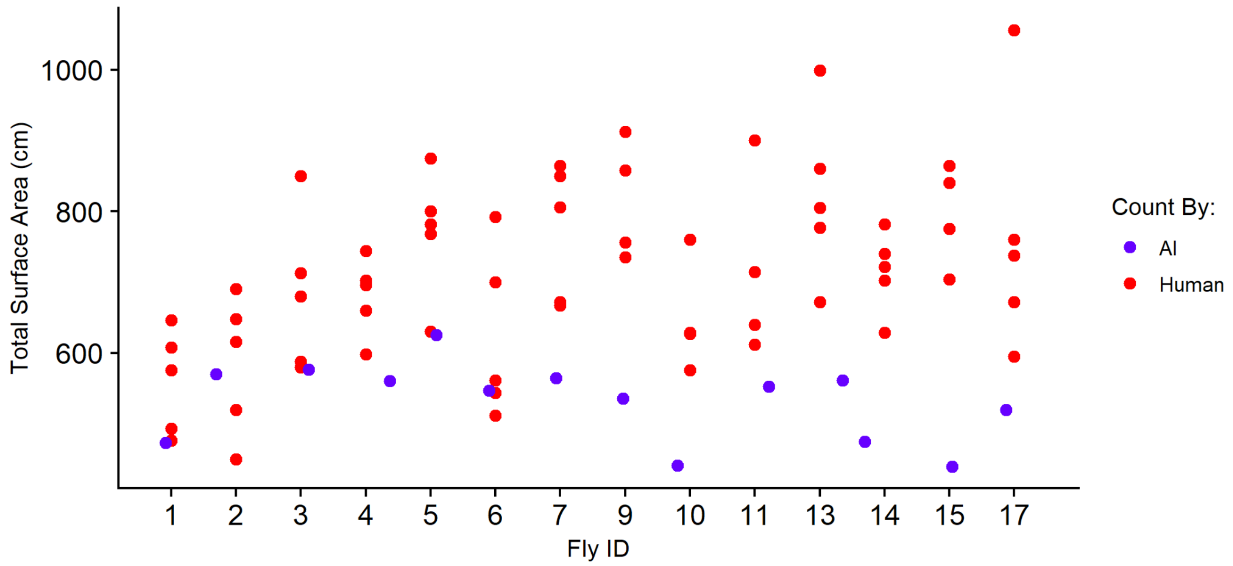


**Figure S5:** Surface area of an object by ID calculated in each frame. The algorithm was trained on: n = 297, 70/30 split, 200 epochs, and plotted against human counts using ImageJ over 5 Frames for each of the IDs. AI counts were calculated by running MLDAAPP and taking the mean of the middle 50% of the values and plotting it according to the IDs. Data extracted using a 60 second sample (1800 frames) of the non-standardized fruit fly environment videos (link in table S1).

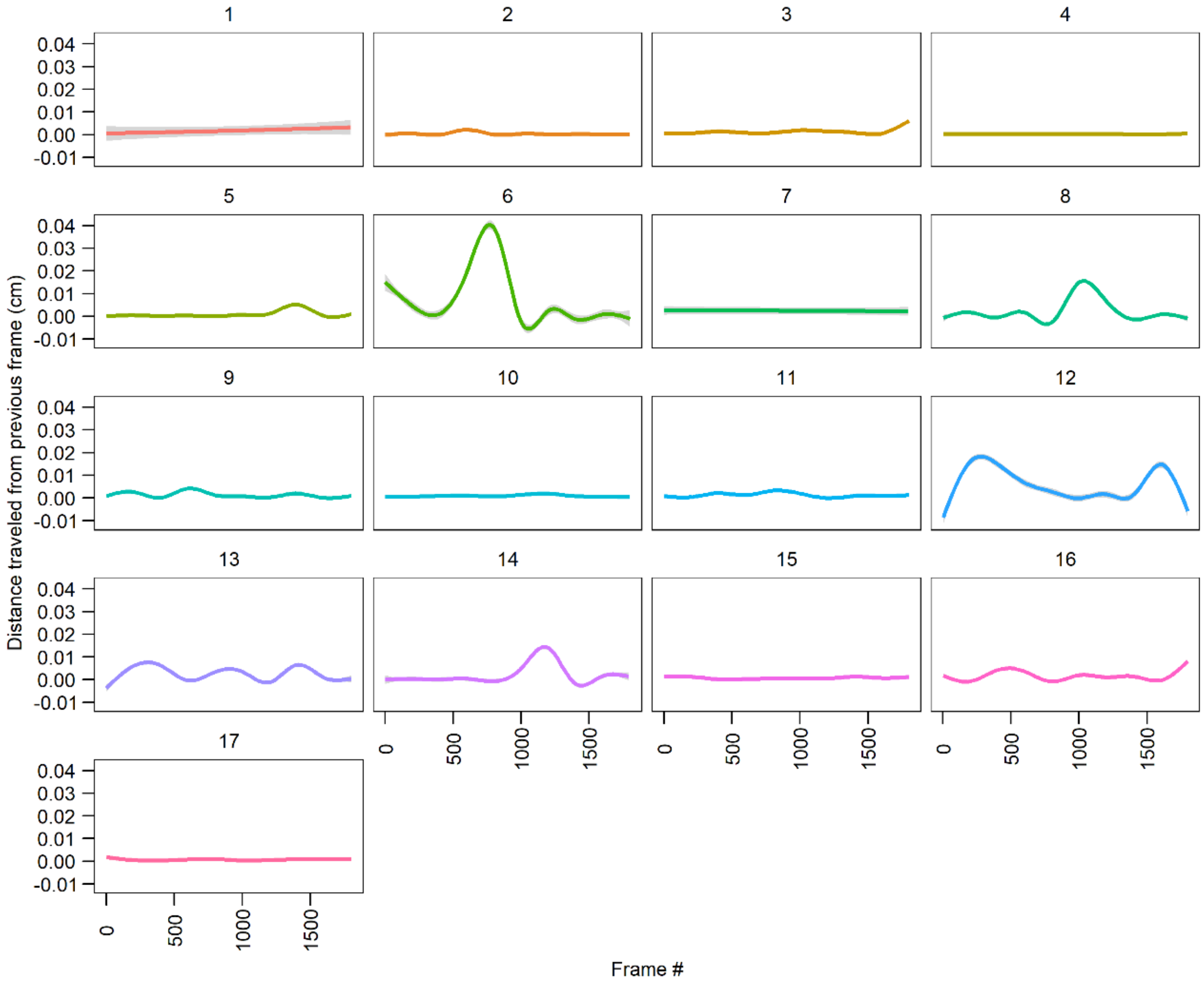


**Figure S6:** Frame by frame movement for each object ID in centimeters. Movement was tracked on a 60 second subset (1800 frames) from the non-standardized fruit fly environment video (link in table S1) using an algorithm based on: n = 297, 70-30% split, 200 epochs.

| Data: | Fecundity | Census | Stand. Flies | Non-Stand. Flies | Daphnia | Rainbow Trout | Humans | Hamsters | Bees |
|---|---|---|---|---|---|---|---|---|---|
| MLDAAPP | A | B | C | D | E | F | G | H | I |
| MLDAAPP vs. Others | | | 1 | 2 | | | | 3 | |

**Table S1:** Links to images and videos used for testing of MLDAAPP. The second row contains annotated videos that directly compare MLDAAPP, IDtracker.ai and TRex.

| Ideal Set-up (No/Little Manual Involvement) | Difficult Set-up (Some/Moderate Manual Involvement) | Unsuitable (Moderate/Large Manual Involvement) |
|---|---|---|
| **Model Choice:**<br>- Yolov8x-seg & Yolov8l-seg<br>**Training set:**<br>- 150+ Unique Images | **Model Choice:**<br>- Yolov8m-seg & Yolov8s-seg<br>**Training set:**<br>- 70-100 Unique Images | **Model Choice:**<br>- Yolov8s-seg & Yolov8n-seg<br>**Training set:**<br>- < 50 Unique Images |
| **Environment:**<br>- Standardized<br>**Object of Interest:**<br>- Clearly Visible | **Environment:**<br>- Standardized or Non-standardized<br>**Object of Interest:**<br>- Clearly Visible or Lightly Obscured | **Environment:**<br>- Non-Standardized<br>**Object of Interest:**<br>- Obscure / Hidden |
| **Image/Video Quality:**<br>- High resolution (Full HD)<br>- High framerate (30+) | **Image/Video Quality:**<br>- Less than 1080p<br>- Less than 30 FPS | **Image/Video Quality:**<br>- Less than 640p<br>- Less than 15 FPS |
| **Tracking Conditions:**<br>- No object overlap<br>- No instantaneous movement | **Tracking Conditions:**<br>- Objects overlap<br>- Rapid Movement | **Tracking Conditions:**<br>- Object overlap<br>- Rapid Movement<br>- Entering/Exiting the Frame |
| **Possible Issues:**<br>- No issues | **Possible Issues:**<br>- Object Loss<br>- ID reassignment<br>- Bad object recall | **Possible Issues:**<br>- Object Loss<br>- ID reassignment<br>- Bad object recall<br>- Low precision |

**Table S2 :** Best Practices for running MLDAAPP to maximize model accuracy and efficiency.
